# BERTrand - peptide:TCR binding prediction using Bidirectional Encoder Representations from Transformers augmented with random TCR pairing

**DOI:** 10.1101/2023.06.12.544613

**Authors:** Alexander Myronov, Giovanni Mazzocco, Paulina Król, Dariusz Plewczynski

## Abstract

**Motivation:** The advent of T cell receptor (TCR) sequencing experiments allowed for a significant increase in the amount of peptide:TCR binding data available and a number of machine learning models appeared in recent years. High-quality prediction models for a fixed epitope sequence are feasible, provided enough known binding TCR sequences are available. However, their performance drops significantly for previously unseen peptides.

**Results:** We prepare the dataset of known peptide:TCR binders and augment it with negative decoys created using healthy donors’ T-cell repertoires. We employ deep learning methods commonly applied in Natural Language Processing (NLP) to train part a peptide:TCR binding model with a degree of cross-peptide generalization (0.66 AUROC). We demonstrate that BERTrand outperforms the published methods when evaluated on peptide sequences not used during model training.

**Availability:** The datasets and the code for model training are available at https://github.com/SFGLab/bertrand

**Supplementary information:** Supplementary data are available at *Bioinformatics* online.

## 1 Introduction

Cytotoxic T-cells play a major role in adaptive immune response in humans. Intracellular proteins are degraded by proteasome into peptides. The antigen processing machinery of the cell allows the presentation of peptides on the cell surface using Major Histocompatibility Complexes (MHC). The focus of this work is MHC class I, which typically presents a peptide of 8 to 11 amino acids long. These peptide-MHC (pMHC) complexes in turn can be recognized and engaged by CD8+ cytotoxic T-cells. Due to negative selection in the thymus and the high degree diversity of the T-cell receptor (TCR) repertoire, T-cells are capable of recognizing a variety of foreign and mutated epitopes (Rudolph *et al*., 2006).

The binding properties of a TCR to a given pMHC is regulated by hypervariable Complementarity Determining Regions (CDRs). For the *α* and *β* chains of the TCR, three such regions exist. The CDR1 and CDR2 of both *α* and *β* chains mostly interact with the MHC complex, while CDR3 is predominantly interacting with the peptide. The CDR3 region is the most variable region of the TCR and is thought to be the major factor in determining the binding preference of the TCR towards its conjugated pMHC (La Gruta *et al*., 2018). While both *α* and *β* chains contribute to the interaction, data suggests that the *β* chain plays the most significant role in target recognition and is significantly more prevalent in the data (Sidhom *et al*., 2021). As this work implies the use of machine learning to infer the interaction between TCRs and pMHC complexes, it can benefit from a high number of observations. Thus we will be focusing solely on the sequence CDR3*β* part of the TCR, which is readily available in multiple datasets.

The MHC is not able to present each and every peptide. Thanks to a high amount of peptide-MHC binding data as well as peptide presentation data from mass spectrometry experiments, researchers were able to produce high accuracy models of peptide-MHC binding and presentation. However, even if the peptide is presented on the surface of the cell, it is still unlikely to be immunogenic. The study by Parkhurst *et al*., 2019 of peptide T-cell immunogenicity for 75 cancer patients has been able to find only 57 CD8+ positive mutations along with over 8000 non-immunogenic ones, which account for less than 1 percent. One of the frontiers of computational immunology research currently is peptide:TCR binding prediction, which represents a key component for understanding T-cell activation. The amount of data for peptide:TCR binding prediction has been growing in recent years, with the popularization of peptide-MHC dextramer production, single-cell TCR sequencing and TCR barcoding. In this work, we compile a collection of peptide:TCR sequence data from a number of databases and publications into a single curated dataset of known TCR binders with their cognate epitope sequences. We augment the dataset with negative decoy examples generated from reference T-cell repertoires.

Machine learning for peptide:TCR binding prediction has been applied before. Glanville *et al*., 2017 and Dash *et al*., 2017 have demonstrated that for a given epitope sequence, TCR binding can be predicted accurately using distance-based methods. In NetTCR2.0 (Montemurro *et al*., 2021), researchers use CNNs to predict peptide:TCR binding. This model performs remarkably well when predicting for a set of peptides from the IEDB database. Another model for peptide:TCR binding predictions is ERGO (Springer *et al*., 2020), where researchers use pre-trained long short-term memory (LSTM) and Autoencoder architectures to predict binding probability and evaluate the performance of the model on several machine learning tasks. Such tasks as single peptide binding, where the predictive power of the model is measured for a single target epitope with different TCRs, or multi-peptide selection, when binding is predicted for a single TCR and a set of possible peptides, have limited applicability in practical scenarios. The goal of this research is to provide immunologists with better tools for *in silico* TCR therapy design. As most of the peptides in the published data originate from viruses, such peptide-centric models can’t be used effectively for cancer neoantigens or tumour-associated antigens. The focus of this work is thus the peptide:TCR pairing task, specifically the case when the model hasn’t previously seen neither the peptide nor the TCR. We believe that high accuracy for this task would bring the most benefit for potential users.

Recent breakthroughs in Natural Language Processing (NLP), such as TAPE (Rao *et al*., 2019) and DNABERT (Ji *et al*., 2021), have prompted many researchers to apply Transformer architectures to solve sequence-based biological problems such as transcription factors prediction, protein-protein interaction prediction, and binding pockets prediction to name a few. One useful feature of models from the NLP domain is the ability to process variable-length sequences of symbols from a fixed alphabet. Another important aspect is the ability to benefit from unsupervised pre-training, which is often an option in bioinformatics. Researchers usually pre-train the language model on large sequence databases, such as UniPROT. In this work, we construct a pre-training set for the peptide:TCR model, which comprises a hypothetical human peptide:TCR repertoire, based on peptides from MHC-I mass spectrometry peptide presentation experiments and synthetic TCRs. After pre-training our model is fine-tuned to predict peptide:TCR binding and was shown to outperform the existing methods in cross-peptide generalization task.

The overall flow of the analysis is demonstrated in Figure 1. In the left part of the figure we show the process of the NLP model pre-training. Synthetic TCRs are paired randomly with the presented peptides to produce a hypothetical peptide:TCR repertoire, which is then used to perform masked language modeling (MLM) pre-training of the BERT neural network. In the right part of the figure the process of generating negative decoy observations is illustrated. Reference TCR sequences from healthy donors’ TCR repertoire sequencing experiments are collected. A machine learning model was used to remove outliers, and then the remaining reference TCRs were clustered together with binding TCRs. TCR sequences that were too similar to any binding TCR were removed, and the rest of the reference TCR clusters were randomly paired with peptides from the binding peptide:TCR set. Pre-trained BERT network was then trained to predict peptide:TCR binding.

**Fig. 1:**
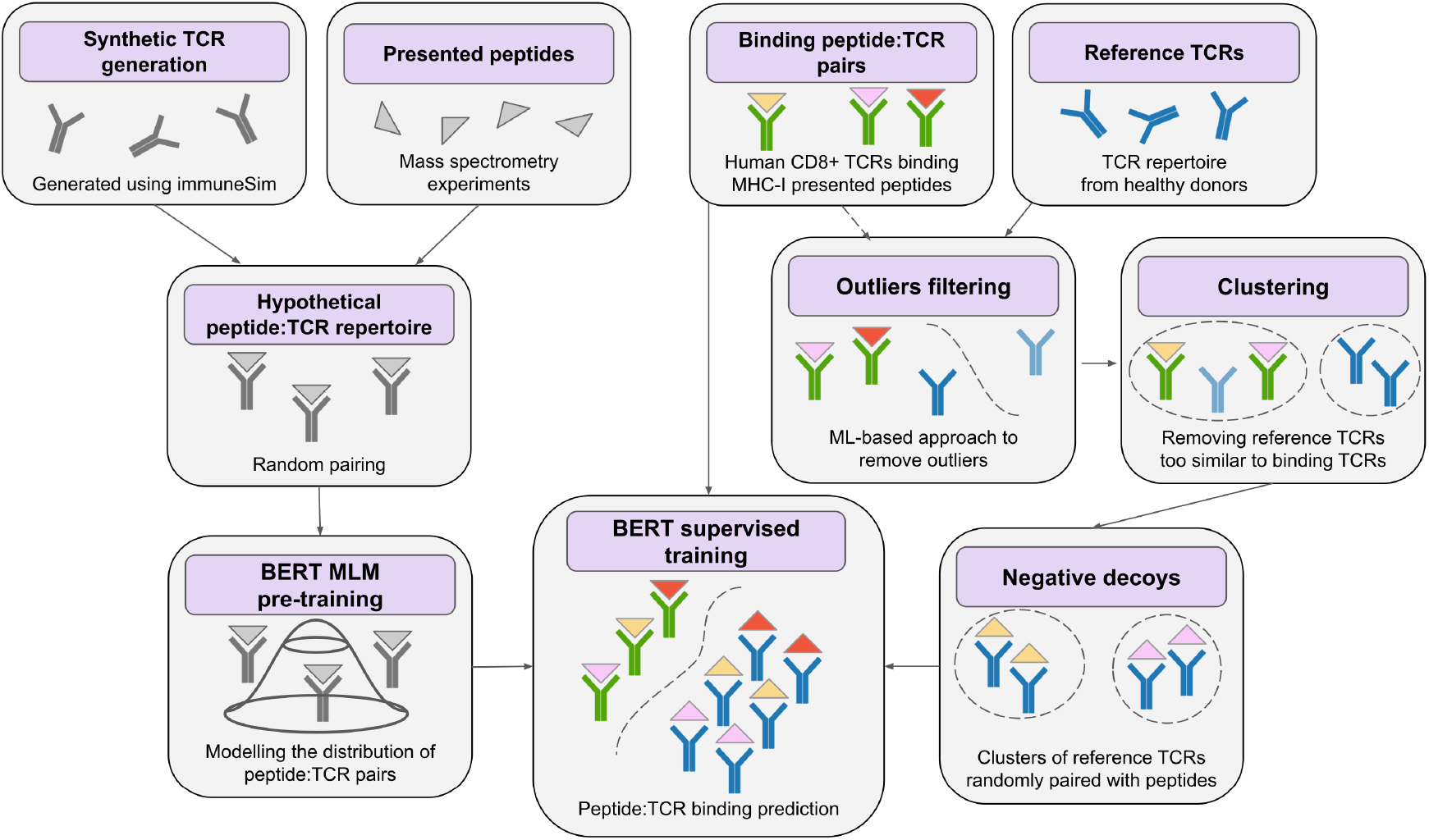
Flow diagram of the analysis. The left part illustrates the creation of the hypothetical peptide:TCR repertoire MLM pre-training. The right part shows the process of negative decoys generation and various filtering steps leading to it. These two paths converge on BERT supervised training for peptide:TCR binding prediction.

## 2 Materials and methods

### 2.1 Data

#### 2.1.1 Data curation

Our data curation process is shown in Figure 2 We collected data containing the aminoacid sequences of binding peptide:TCR pairs from a number of databases and publications (See Table 1). We narrowed down the dataset to human CD8+ T-cells for peptides of 8-11 amino acids long. We only considered CDR3*β* observations, as CDR3*α* annotations are available for only 5% of the data. Over 99% of the CDR3*β* sequences have a length between 10 and 20 amino acids. Another filtering criterion was the requirement of having specific amino acids in the first and last anchor positions in CDR3*β* chains, cysteine (C) and phenylalanine (F) respectively. This way we compiled 33k unique CDR3*β* sequences of T-cells binding with a total of 401 epitope sequences. To compensate for the lack of negative examples, we generated negative decoy observations in a 3-to-1 ratio, using a dataset of reference T-cell repertoires from healthy donors (Oakes *et al*., 2017) and paired it randomly with peptides from the binders dataset.

**Table 1.**
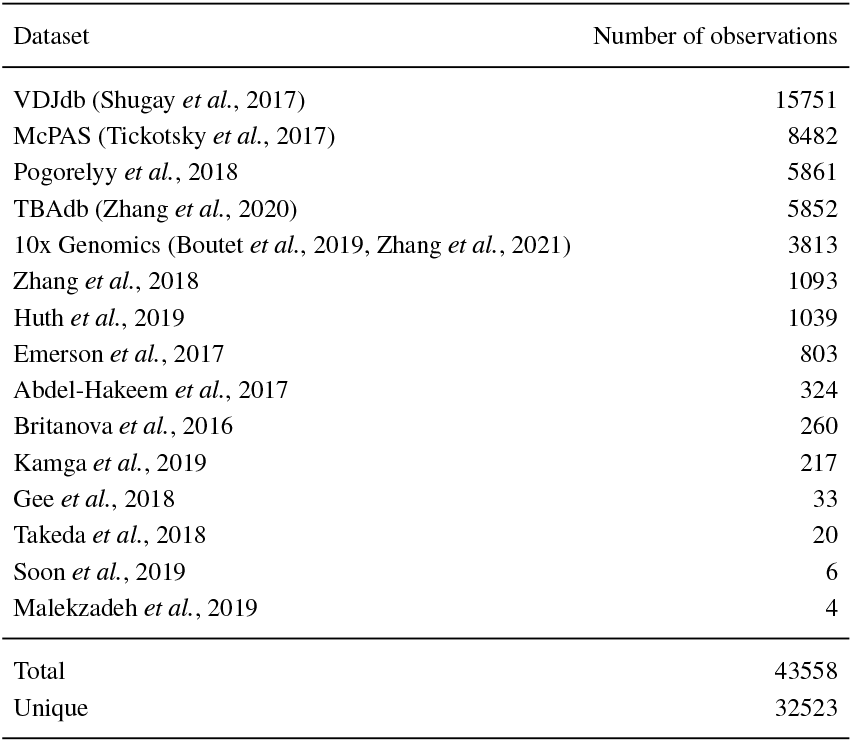
Summary of the binding peptide:TCR dataset. The number of observations is reported after filtering. The curated set of unique peptide:TCR binders is later used to train the model.

| Dataset | Number of observations |
| --- | --- |
| VDJdb (Shugay *et al.*, 2017) | 15751 |
| McPAS (Tickotsky *et al.*, 2017) | 8482 |
| Pogorelyy *et al.*, 2018 | 5861 |
| TBAdb (Zhang *et al.*, 2020) | 5852 |
| 10x Genomics (Boutet *et al.*, 2019, Zhang *et al.*, 2021) | 3813 |
| Zhang *et al.*, 2018 | 1093 |
| Huth *et al.*, 2019 | 1039 |
| Emerson *et al.*, 2017 | 803 |
| Abdel-Hakeem *et al.*, 2017 | 324 |
| Britanova *et al.*, 2016 | 260 |
| Kamga *et al.*, 2019 | 217 |
| Gee *et al.*, 2018 | 33 |
| Takeda *et al.*, 2018 | 20 |
| Soon *et al.*, 2019 | 6 |
| Malekzadeh *et al.*, 2019 | 4 |
| Total | 43558 |
| Unique | 32523 |

**Table 2.** The overview of potential biases that a machine learning algorithm for peptide:TCR prediction can exploit.

| Bias | Mismatch pairing | Reference pairing |
| --- | --- | --- |
| Under-representation of peptides in the dataset | Yes | Yes, but it is mitigated by pre-training to some extent |
| Under-representation of TCRs in the dataset | Yes | No, because additional TCR sequences are used |
| Potential false negatives | Yes | Yes |
| Out-of-distribution easy negative decoys | No | Yes, but they can be filtered to some extent |
| Too homogeneous negative decoys | Yes | No |
| Correlated observations in train and test sets | Yes, because mismatch pairing introduces additional dependencies | No, peptide and TCR clusters are used for decoy generation and cross-validation |

**Fig. 2:**
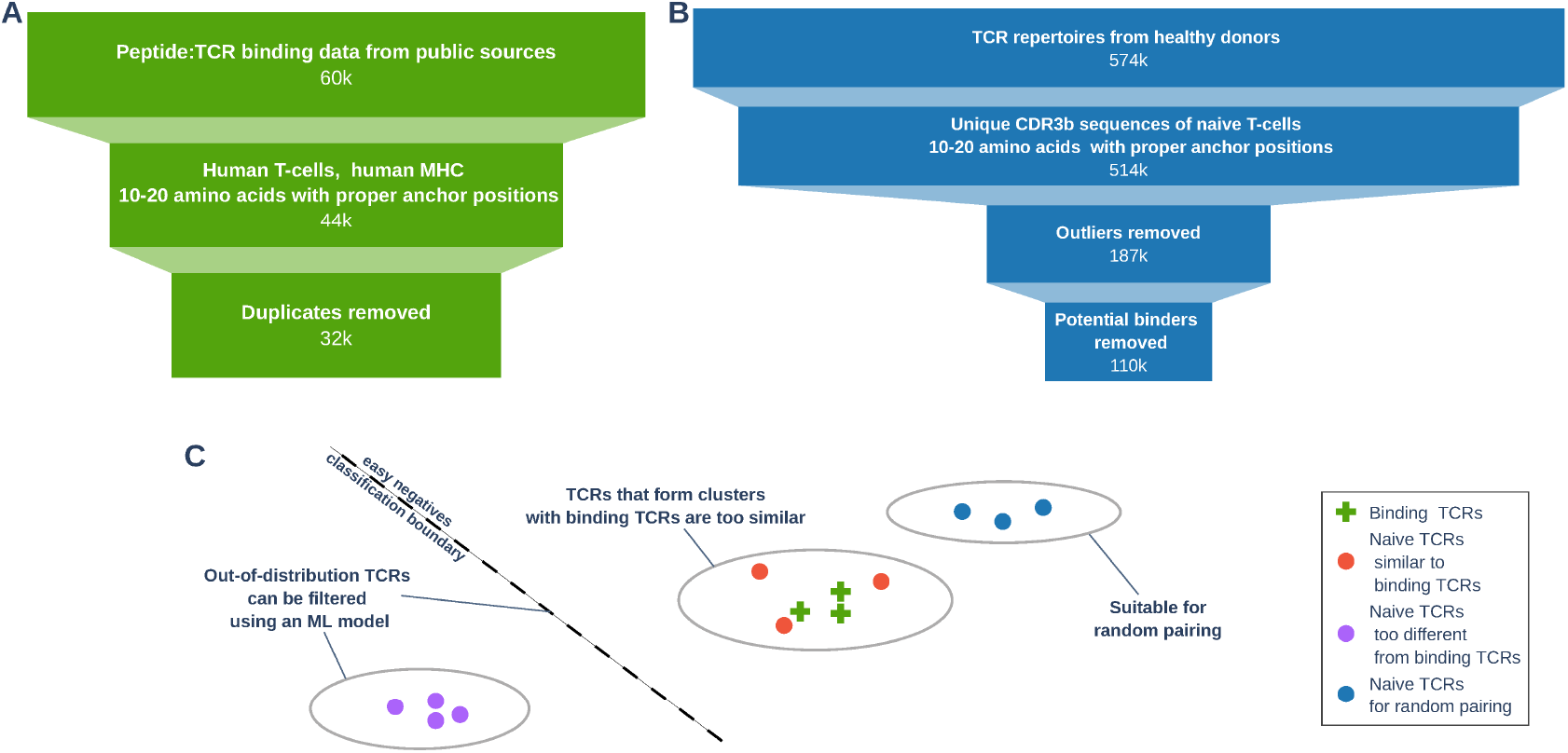
Data curation process. (A) Binding peptide:TCR observations. (B) Reference TCRs for decoy generation. (C) Reference TCRs filtering illustrated.

Cluster analysis of peptides revealed that a number of similar epitope sequences are present, differing by three amino acids at most. For example, MART-1 human melanoma-related antigen (ELAGIGILTV) was extensively studied with minor modifications: EAAGIGILTV, LLLGIGILV, ELAGIGLTV, AAGIGILTV, ALGIGILTV. We argue that using such similar observations in the training and validation set may bias any model trained on peptide sequences and could introduce unwanted overfitting (if the TCR repertoires of two similar peptides are also similar), or unwanted underfitting if the repertoires are too different. Our analysis revealed 261 peptide clusters with a minimum Levenshtein distance of 3. Peptide clusters were used as groups during cross-validation.

#### 2.1.2 Sources of overfitting

Training a machine learning model on a limited set of data (i.e. positive peptide:TCR pairs for 400 peptides only) and with a lack of negative examples forces us to create negative decoy observations, which may easily introduce some bias that an ML model could easily exploit. Randomly pairing peptides and TCRs to create negative decoys is certainly a viable approach for this problem. TCRs are highly cross-reactive and are estimated to bind 10^6^ multiple epitope sequences (Mason, 1998), hence a TCR could potentially bind some other peptide. However, TCRs are also highly specific, as the probability of a specific TCR binding a randomly chosen peptide is estimated to be 10^4^ (Frank, 2020), which is acceptable for a machine learning approach. Besides possible false negative observations produced by random pairing and biases that may arise from experimental conditions, we’ve identified a handful of potential problems for a training setup with negative decoys.

#### 2.1.3 Mismatch pairing

Mismatch pairing is an approach used in Montemurro *et al*., 2021 and Springer *et al*., 2020 to create negative decoy observations by randomly pairing a peptide with a TCRs from a different peptide:TCR pair. Mismatch pairing approach guarantees that the TCR distribution of the negatives will match the positive one. The number of different TCRs in a human body is estimated to be around 10^10^ (Lythe *et al*., 2016), and the number of MHC-I-presented peptides on human cells is around 10^4^ (Mester *et al*., 2011), thus mismatch pairing is largely under-representing both peptides and TCRs. Moreover, due to publication bias, different potential immunogenicity of viral and cancer peptides in humans, the distribution of T-cell clonotype size in humans (Bolkhovskaya *et al*., 2014) and other factors, the number of unique TCR observations per peptide has a power-law decay distribution (See Figure 1 in Supplementary data). In practice this means that random peptide:TCR pairing will produce negative decoys, where over 61% of TCR sequences come from the 5 most popular peptides in the dataset. The repertoires of these peptides are too homogeneous and a model might learn to identify those as negatives. Mismatch pairing also produces correlated observations, thus during cross-validation there would be a considerable amount of examples in the training set sharing peptide or TCR sequence with pairs in the test set. These recurring peptides and TCRs may be exploited by the model if not removed.

#### 2.1.4 Reference pairing

Our approach to negative decoys generation was designed to overcome some of the aforementioned biases. We collected around 560k TCR CDR3*β* sequences from repertoires of 3 healthy donors from Oakes *et al*., 2017 and randomly paired them with peptides from the binding dataset CDR3*β* sequences. Reference TCRs represent a much larger region of the TCR theoretical distribution compared to binding TCRs, although both TCRs and peptides remain still under-represented. Reference TCRs might introduce CDR3*β* sequences that are out-of-distribution relative to the binding TCRs and thus might be easy targets for a neural network. To address this issue, we filtered the sequences using a ML-based approach, see Outliers filtering section in Supplementary data for a detailed description.

To avoid correlated observations, we performed a TCR clustering analysis of the binding TCR sequences together with reference TCRs: agglomerative clustering was used with Levenshtein distance and complete linkage with distance cut-off equal to 3. Clustering naturally produced 3 types of clusters:

1. clusters with only TCRs from binding peptide:TCR pairs
2. mixed clusters positive and reference TCRs
3. clusters with only reference TCRs

During decoy generation, we rejected reference TCRs from mixed clusters, as they have a much higher probability of being false negatives, because their sequences are very similar to those of the binding TCRs. TCRs from reference-only clusters were randomly paired with a single peptide from the pool of 401 available peptides in the binding dataset in a 3-to-1 ratio to the number of positive observations for a given peptide. In a cluster of reference TCRs, every TCR is paired with the same peptide, which limits the generation of positive and negative observations with the same peptide sequence and very similar TCR sequences. Such observations would in fact be quite useless, as they should be filtered from the training set if they appear in the test set during cross-validation, otherwise the model would be biased towards predicting the negative class. As the decoy generation is random, the dataset was replicated three times for different seeds. TCR clusters were also used as groups during cross-validation.

### 2.2 Model

For this problem, we applied the Bidirectional Encoder Representations from Transformers (BERT) artificial neural network (Devlin *et al*., 2019) from *transformers* python package (Wolf *et al*., 2019). The model was initially pre-trained to perform a masked language modeling (MLM) task on a hypothetical TCR repertoire and then fine-tuned for actual peptide:TCR classification.

#### 2.2.1 Model architecture

The architecture of BERT is illustrated in Figure 3. Initially, peptides and TCRs need to be represented as a sequence of tokens. The token vocabulary consists of 20 amino acids and 5 additional special tokens: *CLS* is used to indicate the starting position of the sequence, *SEP* to indicate the end of the sequence, *MASK* for masked language modeling, *PAD* for padding, and *UNK* for non-standard amino acids. Each peptide and TCR are concatenated into a single sequence of tokens with an additional *CLS* token at the beginning and two *SEP* tokens between sequences and at the end of the sequence. The sequence IDs indicating the absolute position of the token and the token type IDs indicating whether a token belongs to the peptide or to the TCR are generated as well. Token, position and type IDs are embedded and added creating the sequence of input embeddings for each token. The input is then passed to a BERT network with 8 transformer blocks. BERT produces an output embedding for each token, which later on is passed to either a token classification head and to a sequence classification head during pre-training and fine-tuning, respectively. Below is the description of the BERT embedding procedure.

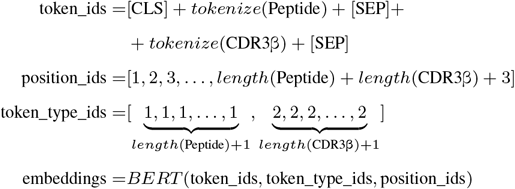

**Fig. 3:**
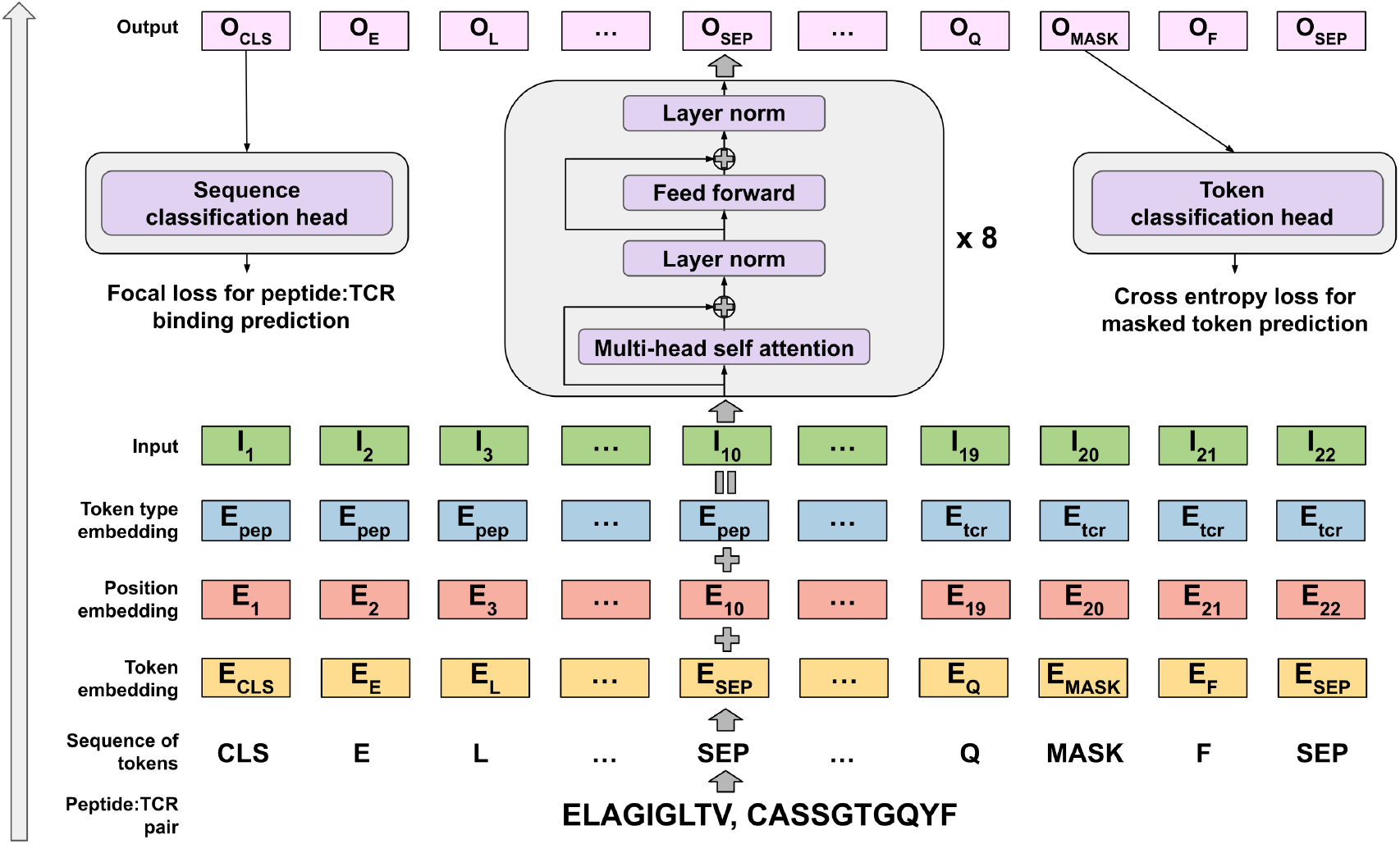
BERT architecture (bottom to top): First, peptide and TCR are tokenized. Then, each token is embedded in a 512-dimensional token space (yellow). The absolute position (red) and token type (blue) - peptide or TCR - are encoded using separate embeddings. All 3 embeddings are added to form the input (green). 8 Transformer blocks (violet) process the input to produce the output for each token (pink). During MLM pre-training, the token classification head (right) is trained to predict the true token 15% randomly masked tokens. During sequence classification training (left), the sequence classification head takes output of the CLS token and predicts binding for a peptide:TCR pair

### 2.2.2 Pre-training

The large (over 2.5M) number of trainable parameters in BERT is compensated by the unsupervised pre-training strategy. For this purpose, we created the hypothetical peptide:TCR repertoire. 11M CDR3*β* sequences were obtained by simulating V(D)J recombination using immuneSim (Weber *et al*., 2020). 150k peptides were acquired from the results of mass spectrometry peptide sequencing experiments (Abelin *et al*., 2017, Di Marco *et al*., 2017, Faridi *et al*., 2018, Sarkizova *et al*., 2020). We randomly paired the MHC-I-presented peptides with simulated TCRs and pre-trained a BERT neural network using masked language modeling (MLM). 15% of randomly chosen amino acids in the input sequence were masked and the network was trained to predict the masked amino acid. The weights from this stage were used as a starting point for the supervised training task. The effect of MLM pre-training can be seen in Figure 2 in Supplementary data. The network was pre-trained for 100 epochs on a dataset of 4.5M peptide:TCR pairs. See Hyperparameters section in Supplementary data for more details about model training. Below is the description of the BERT MLM procedure.

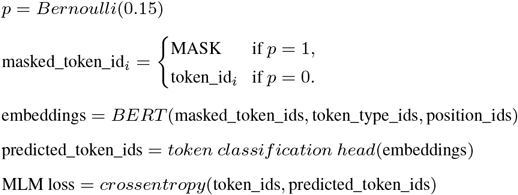

### 2.2.3 Fine-tuning

A separate sequence classification head was trained to classify binding peptide:TCR pairs and negative decoys. The hidden representation of the first token in the sequence was passed into a feed-forward layer combined with a softmax layer output. Focal loss with γ=3 and α=0.25 was used. Below is the description of the BERT fine-tuning procedure.

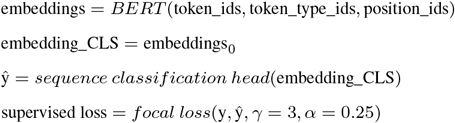

### 2.3 Benchmarks and evaluation

Existing approaches to peptide:TCR binding predictions based on peptide and CDR3*β* sequences were evaluated alongside our model, namely NetTCR2.0 and ERGO. NetTCR was published by Montemurro *et al*., 2021, who used a convolutional neural network to predict peptide:TCR binding for A02:01-restricted peptides. NetTCR performs very well on peptide sequences that the model has already seen. ERGO is the algorithm published by Springer *et al*., 2020, which uses a pre-trained Long short-term memory neural network (LSTM) to produce a model that performs well for both known and unknown peptides and TCRs.

To test the generalization power of our model, we performed a repeated cross-validation grouped by peptide and TCR clusters, to avoid correlated observations in the training and testing set. For each training episode the dataset was split into training, early stopping and test sets. 14 peptide clusters were chosen for the test set. Any TCR present in clusters containing test set observations was removed from both training and early stopping sets. All models were trained using the same dataset splits to ensure fairness in terms of data availability. The training, early stopping and test sets were restricted to viral peptides and the final quality of the predictions was measured using the AUROC metric on an independent set of cancer peptides.

The model and the benchmarks were trained for 25 epochs. BERTrand and ERGO use early stopping on a separate set, however NetTCR2.0 uses training loss for early stopping. The metric used for model validation was AUROC, which was computed separately for each peptide and then averaged. Using this averaging procedure, we limit the bias which originates from the high number of observations for some peptides and from the differences between peptide repertoires. We believe that such a measure is preferable to AUROC computed across all the observations, as the latter might likely lead to inflated results. Every model was also evaluated using an independent test set of cancer peptides.

## 3 Results and discussion

We performed 21 rounds of cross-validation, repeated 3 times with different random pairings, and we evaluated the average per-peptide AUROC for each model. BERTrand converged to the optimal solution at around 5 epochs, with further training only introducing overfitting (See Figure 3 in Supplementary data). The cross-validation results shown in Figure 4 indicates that our model can achieve better predictive performance than the state-of-the-art models shown in the benchmarks when tested in multiple scenarios. While the average AUROC indicates a model that is definitely better than random, the high variability in AUROC between peptides is definitely a concern. Out of 76 cancer peptides the BERTrand achieved over 0.55 AUROC for 58 of them. While the model may be not optimally suited to perform prediction tasks on single peptide targets, it is applicable to groups of peptide targets in a practical *in silico* TCR prioritization scenario. With a large enough set of previously unseen peptide targets, BERTrand can generate predictions that prioritize binding peptide:TCR pairs yielding an expected AUROC of 0.66.

**Fig. 4:**
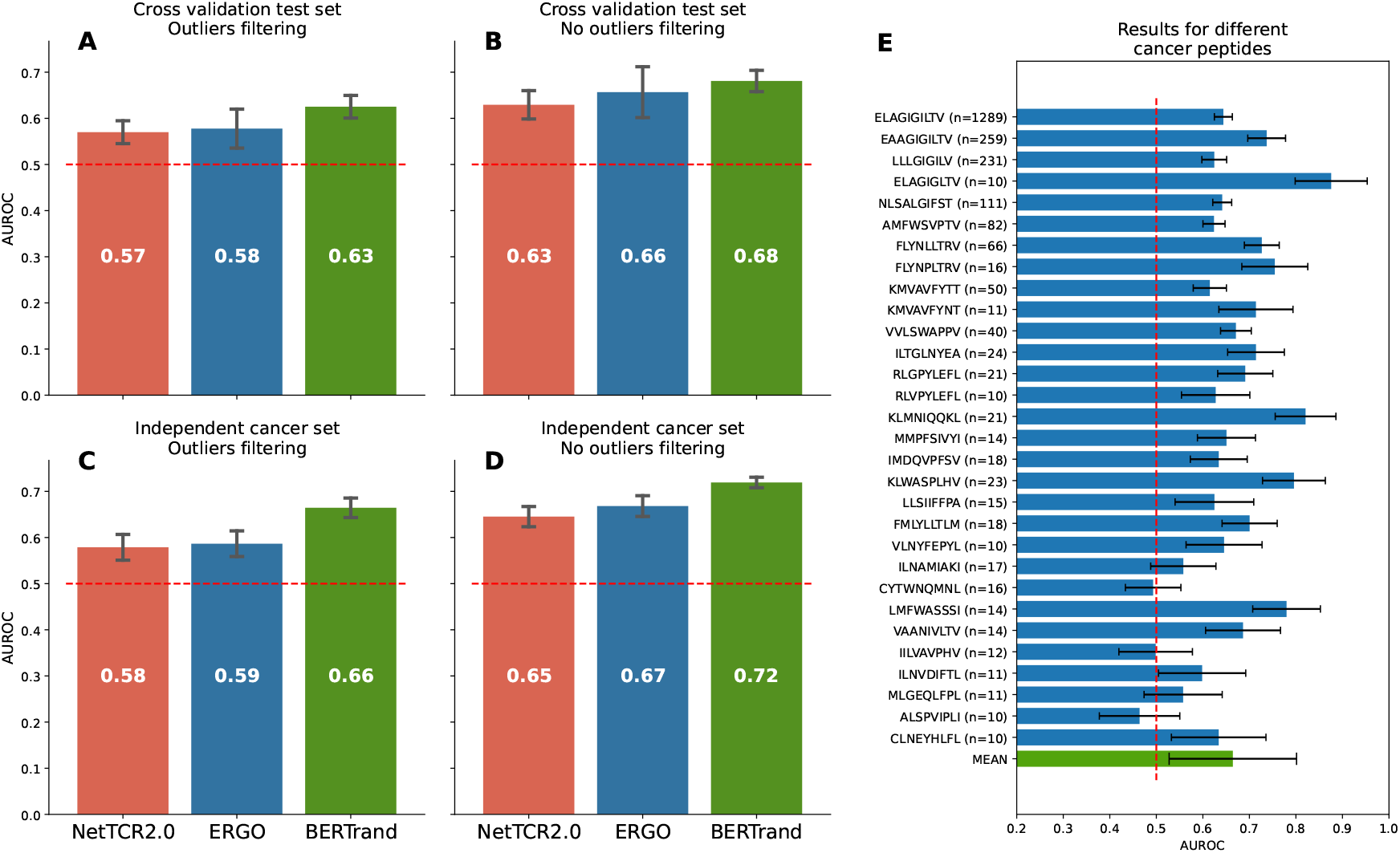
Benchmarks. (A) Results from cross-validation. 14 peptide clusters were sampled at the beginning of each training episode, the process was repeated 21 times for 3 random pairings. (B) Results from cross validation without easy negatives filtering demonstrate the potential metrics inflation due to outliers. (C) Results on an independent cancer set of observations - BERTrand demonstrates better performance that two other models. (D) Results on cancer peptides without easy negatives filtering are inflated and again demonstrate the importance of outliers filtering. (E) Results for individual peptides in the independent cancer set. Note that the confidence interval of the mean AUROC is wider as it includes high per-peptide variation.

We believe that the biggest challenge in peptide:TCR binding prediction is peptide bias due to low diversity of the peptides available. It is obvious that neither the peptide space (i.e. all possible peptides to be bound by TCRs) nor the TCR space (all possible TCRs) is sufficiently explored in the published data. NetTCR2.0 uses convolutional neural networks to predict the binding, which are very susceptible to overfitting when an obvious bias, such as a very popular peptide in the training dataset, is present. ERGO overcomes the TCR diversity problem by applying unsupervised TCR pre-training, but the peptide bias remains unaddressed. Joint peptide:TCR pre-training strategy might help to address this bias, as we have shown in this work. Language-based models can be successfully applied to the peptide:TCR binding problem outperforming other state-of-the-art methods, as highlighted in the results of our tests.

## 4 Conclusions

Cross-peptide generalization in peptide:TCR binding prediction remains a hard problem. However, results presented in this work demonstrate that peptide:TCR binding is predictable beyond known peptide targets. We believe the biggest obstacle in this field is data availability. Recent advances in single-cell TCR sequencing allowed for producing more experimental data, so we hope this work will encourage future peptide:TCR binding experiments and in the end, allow predictive models to become very useful for researchers. Potential applications of the peptide:TCR prediction model include the design of off-the-shelf TCR-based therapies for cancer (e.g. TCR-engineered T-cell therapies, TCR-mimicking antibodies, TCR bispecific antibodies), the development of de-immunization strategies for autoimmune diseases, and the selection of optimal candidates for antiviral vaccine design. Even a model with a limited predictive power can already represent a very useful tool for the optimization of *in vitro* experiments (e.g. reducing the experimental time and costs associated with the typical experimental testing of a large number of putative non-prioritized TCR candidates.

An important limitation of this work is the lack of CDR3*α* due to low availability of those sequences in databases. Although CDR3*α* sequence has been reported to be important for peptide:TCR binding prediction (Sidhom *et al*., 2021), it is only available for less than 5% of observations. Due to the low MHC diversity in the data, the aspects of the interactions between the MHC and the TCR were also omitted. However, we are confident that the growing collective effort we are witnessing in this field will eventually lead to populating databases with large amounts of fully annotated data. We believe this will open new doors for a holistic solution to the pMHC: TCR binding problem, with profound consequences for the development of immune-related therapies.

## Acknowledgements

We would like to thank our fellows at the Laboratory of Bioinformatics and Computational Genomics at the Warsaw University of Technology for providing valuable suggestions and being a great audience.

## Funding

This work was supported by the PhD programme “Doktoraty Wdzrozeniowe” of the Polish Ministry of Education and the research grant “Stworzenie innowacyjnej, opartej na sztucznej inteligencji technologii TCRact w celu wprowadzenia na rynek nowej us ł ugi polegającej na projektowaniu in silico receptorów limfocytów T (TCR) do wykorzystania w immunoterapiach nowotworów” (grant number POIR.01.01.01-00-0019/20) by the The National Centre for Research and Development in Poland. Research was co-funded by Warsaw University of Technology within the Excellence Initiative: Research University (IDUB) programme. This work has been co-supported by Polish National Science Centre (2019/35/O/ST6/02484). Computations were performed thanks to the Laboratory of Bioinformatics and Computational Genomics, Faculty of Mathematics and Information Science, Warsaw University of Technology using Artificial Intelligence HPC platform financed by Polish Ministry of Science and Higher Education (decision no. 7054/IA/SP/2020 of 2020-08-28).

## Notes

https://github.com/SFGLab/bertrand

### Competing Interest Statement

The authors have declared no competing interest.

